# A single Ho-induced double-strand break at the *MAT* locus is lethal in *Candida glabrata*

**DOI:** 10.1101/2020.01.27.920876

**Authors:** Laetitia Maroc, Youfang Zhou-Li, Stéphanie Boisnard, Cécile Fairhead

**Affiliations:** Université Paris-Saclay, INRAE, CNRS, AgroParisTech, GQE - Le Moulon, 91190, Gif-sur-Yvette, France; Université Paris-Saclay, CEA, CNRS, Institute for Integrative Biology of the Cell (I2BC), 91198, Gif-sur-Yvette, France

**Keywords:** Mating-type switching, Ho, Homologous recombination, yeast, CRISPR-Cas9

## Abstract

Mating-type switching is a complex mechanism that promotes sexual reproduction in Ascomycotina. In the model species *Saccharomyces cerevisiae*, mating-type switching is initiated by the Ho endonuclease that performs a site-specific double-strand break (DSB) at *MAT*, repaired by homologous recombination (HR) using one of the two silent mating type cassettes, *HMLalpha* and *HMRa*. The reasons why all the elements of the mating-type switching system have been conserved in some Ascomycotina, that do not show a sexual cycle nor mating-type switching, remain unknown. To gain insight on this phenomenon, we used the opportunistic pathogenic yeast *Candida glabrata*, phylogenetically close to *S. cerevisiae,* and for which no spontaneous and efficient mating-type switching has been observed. We have previously shown that expression of *S. cerevisiae*’s *HO* gene triggers mating-type switching in *C. glabrata*, but this leads to massive cell death. In addition, we unexpectedly found, that not only *MAT* but also *HML* was cut in this species, suggesting the formation of multiple chromosomal DSBs upon *HO* induction.

We now report that *HMR* is also cut by *S. cerevisiae*’s Ho in wild-type strains of *C. glabrata.* To understand the link between mating-type switching and cell death in *C. glabrata*, we constructed strains mutated precisely at the Ho recognition sites. By mimicking *S. cerevisiae*’s situation, in which *HML* and *HMR* are protected from the cut, we unexpectedly find that one DSB at *MAT* is sufficient to induce cell death. We demonstrate that mating-type switching in *C. glabrata* can be triggered using CRISPR-Cas9, without high lethality. We also show that switching is Rad51-dependent, as in *S. cerevisiae* but that donor preference is not conserved in *C. glabrata.* Altogether, these results suggest that a DSB at *MAT* can be repaired by HR in *C. glabrata*, but that it is prevented by *S. cerevisiae*’s Ho.

**Author summary:** Mating-type switching is one of the strategies developed by fungi to promote crossing, sexual reproduction and propagation. This mechanism enables one haploid cell to give rise to a cell of the opposite mating-type so that they can mate together. It has been extensively studied in the model yeast *S. cerevisiae* in which it relies on a programmed double-strand break performed by the Ho endonuclease at the *MAT* locus which encodes the key regulators of sexual identity. Little is known about why the mating-type switching components have been conserved in species like *C.glabrata,* in which neither sexual reproduction nor mating-type switching is observed. We have previously shown that mating-type switching can be triggered, in *C. glabrata*, by expression of the *HO* gene from *S. cerevisiae* but this leads to massive cell death. We report here evidence toward a degeneration of the mating-type switching system in *C. glabrata*. We demonstrate that the DSB at *MAT* is only lethal when the Ho endonuclease performs the break, a situation unique to *C. glabrata.* Finally, we show that mating-type switching in *C. glabrata* can be triggered by CRISPR-Cas9 and without any high lethality.

## Introduction

In eukaryotes, sexual reproduction is a nearly ubiquitous feature and implies fundamental conserved processes such as gamete fusion, zygote formation and meiosis (1). Sexual reproduction leads to genetic recombination between organisms and thus enables them to purge their genomes from deleterious mutations, as well as to increase their genetic diversity. It is in the fungal kingdom that the greatest diversity of sexual reproduction is found (1). Particularly, sexual reproduction in fungal human pathogens exhibits a considerable plasticity between species (2) (3). While many were thought to be asexual, several atypical sexual or parasexual cycles have been discovered. It has been shown that the yeast *Candida albicans* can perform a parasexual cycle by mating of two diploid cells, forming a tetraploid one, that can undergo chromosome loss (4). The more distant filamentous opportunistic pathogen, *Aspergillus fumigatus* exhibits a sexual cycle but only mates after spending 6-12 months in the dark (5). Altogether, this suggests that, in most fungi, performing genetic exchange is capital, even in well-adapted human pathogens.

In fungi, sexual reproduction can occur through three mechanisms (1): heterothallism (requiring two compatible partners for mating to occur), homothallism (self-fertility), and pseudo-homothallism (where a single individual can go through a complete sexual cycle but mating only occurs between two compatible partners). Pseudo-homothallism has mainly been described in ascomycete yeasts and occurs through a programmed differentiation process called mating-type switching (6). This mechanism enables one haploid cell to give rise to a cell of the opposite mating-type so that they can mate together. It implies a genomic DNA rearrangement of the MATing-type locus (*MAT*, encoding the key regulators of sexual identity) and species have evolved very different molecular pathways for the same aim. In the fission yeast *Schizosacharomyces pombe*, an imprint at *mat1* is introduced that leads to a DSB during DNA replication (7, 8). Repair occurs with one of the two silent copies of *mat1, mat2* and *mat3*. In the ascomycete *Kluyveromyces lactis,* mating-type switching involves a DSB at *MAT* but it is performed by two specific nucleases depending on the mating-type of the cell (9). Mating-type switching has been extensively described particularly in the model yeast *S. cerevisiae* and has notably allowed a better understanding of cell identity, DSB repair and silencing mechanisms (10).

In *S. cerevisiae*, haploid cells can be of either mating-type, *MATalpha* or *MATa*, which encodes “alpha” or “a” information, respectively, at the Y sequence of the *MAT* locus (11). Mating-type switching relies on a programmed DSB at the *MAT* locus performed by the Ho endonuclease at its 24-bp recognition site. DSBs are highly toxic DNA lesions, and thus have to be efficiently repaired to ensure cell viability. This can be achieved through two major pathways, non-homologous end-joining (NHEJ) and homologous recombination (HR) in the presence of a repair template. The DSB at *MAT* is repaired ∼90% of the time by HR (10), probably because of efficient resection of the DSB that has been shown to prevent NHEJ (12). The Ho cut at the *MAT* locus generates 4 bp, 3′-overhanging ends and its repair implies the following steps. The DSB ends are processed by several 5′ to 3′ exonucleases to create long 3′-ended tails (13). Single-strand tails are then converted to Rad51-coated nucleoprotein filaments, which search for homology and promote homologous template invasion (10). Once the homologous donor is found, the *MAT* locus is repaired by gene conversion. The homologous donor is one of the two silent loci located on the same chromosome as *MAT*: *HML* carrying the “alpha” information or *HMR* carrying the “a” information. The “alpha” or “a” sequence from *HML* or *HMR* respectively, replaces the original Y *MAT* sequence whereas *HML* and *HMR* remain unchanged. Despite the fact that *HML* and *HMR* contain the Ho recognition site, both are resistant to Ho cleavage, being located in heterochromatic regions (14). In *S. cerevisiae*, a “donor preference” mechanism ensures an efficient mating-type switching at *MAT* by promoting the use of the silent locus from the opposite mating-type (*MATa* is preferentially repaired by *HML* and *MATalpha* by *HMR*). This donor preference depends on both the “a” or “alpha” information at the *MAT* locus and the presence of a specific sequence, the recombination enhancer (RE), located between *HML* and *MAT* (15).

*C. glabrata* is an opportunistic pathogenic yeast, phylogenetically close to *S. cerevisiae* (16). Its genome has retained the three-locus system, with homologs of *HML*, *MATa/alpha*, and *HMR*, called <u>M</u>ating-Type <u>L</u>ike (*MTL*) loci (17). The three loci display a structure comparable to *S. cerevisiae’s,* the main difference being that *HMR* is located on a different chromosome from *HML* and *MAT* (17). Despite these similarities, added to the fact that both *MATa* and *MATalpha* cells are isolated and that they maintain mating-type identity (17) (18) (19), *C. glabrata* is unable to switch mating-type spontaneously at an efficient level, even though rare signs of mating-type switching are observed in culture (20) and in populations (21). We have previously shown that the expression of the *HO* gene from *S. cerevisiae* can trigger mating-type switching in *C. glabrata*, and that over 99 % of *C. glabrata* cells are unable to survive to the expression of Ho (22). Conversely, we did not observe mating-type switching after the expression of the *HO* gene from *C. glabrata* in *S. cerevisiae.* By analysing surviving colonies of *C. glabrata* cut by *S. cerevisiae*’s Ho, we had also observed gene conversion events at the *HML* locus, revealing that, contrary to *S. cerevisiae*, *HML* is not protected from the Ho cut. We suggested that the lethality was due to multiple chromosomal DSBs, which would prevent homologous recombination with an intact template in most cells.

In this work, we explore the link between mating-type switching and lethality. For this purpose, we constructed a series of inconvertible **(inc)** strains, mutated precisely at the Ho recognition site, allowing us to control the number and position of DNA breaks during Ho induction, as well as to track which donor sequence is used as template. We analyze two aspects: viability, that reflects both the efficiency of the cut and the success of repair; and molecular analysis of repaired loci, in order to reveal which repair pathways were used. We now show that *HMR* is also cut by Ho in wild-type strains of *C. glabrata.* In addition, by mimicking *S. cerevisiae*’s situation, in which *HML* and *HMR* are protected from the cut, we unexpectedly find that one DSB at the *MAT* locus is sufficient to induce cell death. The use of the CRISPR-Cas9 technology enables us, not only to show that mating-type switching can be induced independently of the Ho protein in *C. glabrata*, but also, that it can be induced without any high lethality.

## Results

### <u>HMR</u> is cut by Ho in <u>C. glabrata</u> and the subsequent mating type switching relies on homologous recombination

As previously described, expression of *S. cerevisiae*’s *HO* gene in wild-type strains of *C. glabrata*, leads to the death of about 99.9 % of cells and we found that both *MAT* and *HML* are efficiently cut (22). We further analyzed surviving colonies of HM100 (*HMLalpha MATalpha HMRa*) by determining the mating-type at each *MTL* locus by PCR and we found that some present switching at *HMR,* indicative of cutting (not shown). Ho-induced lethality in *C. glabrata* could be due to the concomitant induction of multiple DSBs, in contrast to the situation in *S. cerevisiae* where *HML* and *HMR* are protected from the cut, as we hypothesized in our previous work (22).

In order to identify the repair pathway involved in mating-type switching, we inactivated *RAD51* (CAGL0I05544g) in strain HM100 (Table 1).

**Table 1.** Strains used in this work.

| ***C. glabrata* strains with wild-type Ho sites** |  |  |  |
| --- | --- | --- | --- |
| **Strains** | **Parent** | **Genotype** | **Reference** |
| CBS138 |  | *HMLalpha MATalpha HMRA* | (23) |
| BG2 |  | *HMLalpha MATa HMRA* | (24) |
| BG14 | BG2 | *HMLalpha MATa HMRA ura3Δ::Tn903 G418<sup>R</sup>* | (24) |
| BG87 | BG14 | *HMLalpha MATa HMRA ura3::Neo<sup>R</sup> his3Δ* | (25) |
| HM100 | CBS138 | *HMLalpha MATalpha HMRA ura3Δ::KANMX* | (17) |
| HM100 *Δrad51* | HM100 | *HMLalpha MATalpha HMRA Δrad51 ura3Δ::KANMX* | This work. |
| ***C. glabrata* strains with mutated Ho sites** |  |  |  |
| YL01 | HM100 | *HMLalpha MATalpha HMRA-inc ura3Δ::KANMX* | This work. |
| YL02 | HM100 | *HMLalpha-inc MATalpha HMRA ura3Δ::KANMX* | This work. |
| YL03-MATalpha | YL02 | *HMLalpha-inc MATalpha HMRA-inc ura3Δ::KANMX* | This work. |
| YL03-MATa | YL03-MATalpha | *HMLalpha-inc MATa HMRA-inc ura3Δ::KANMX* | This work. |
| YL04 | YL07 | *HMLalpha MATalpha-inc HMRA ura3Δ::KANMX* | This work. |
| YL05 | YL09 | *HMLalpha MATa-inc HMRA-inc ura3Δ::KANMX* | This work. |
| YL07 | YL02 | *HMLalpha-inc MATalpha-inc HMRA ura3Δ::KANMX* | This work. |
| YL09 | YL01 | *HMLa-inc MATa-inc HMRA-inc; ura3Δ::KANMX* | This work. |
| YL10 | YL07 | *HMLalpha-inc MATalpha-inc HMRalpha-inc ura3Δ::KANMX* | This work. |
| SL09 | BG87 | *HMLa-inc MATa-inc HMRA-inc ura3::Neo<sup>R</sup> his3Δ* | This work. |
| ***C. glabrata* strains with mutated Ho sites and/or deletion of *HML* or *HMR*** |  |  |  |
| CGM460 | BG14 | *Δhml MATalpha HMRA ura3Δ::Tn903 G418<sup>R</sup>* | (18) |
| SL01 | BG87 | *HMLalpha MATa Δhmr ura3::Neo<sup>R</sup> his3Δ* | This work. |
| SL-CG8 | SL01 | *HMLalpha-inc MATa Δhmr ura3::Neo<sup>R</sup> his3Δ* | This work. |
| SL-CG9 | CGM460 | *Δhml MATa HMRalpha-inc ura3Δ::Tn903 G418<sup>R</sup>* | This work. |

**Fig 1.**
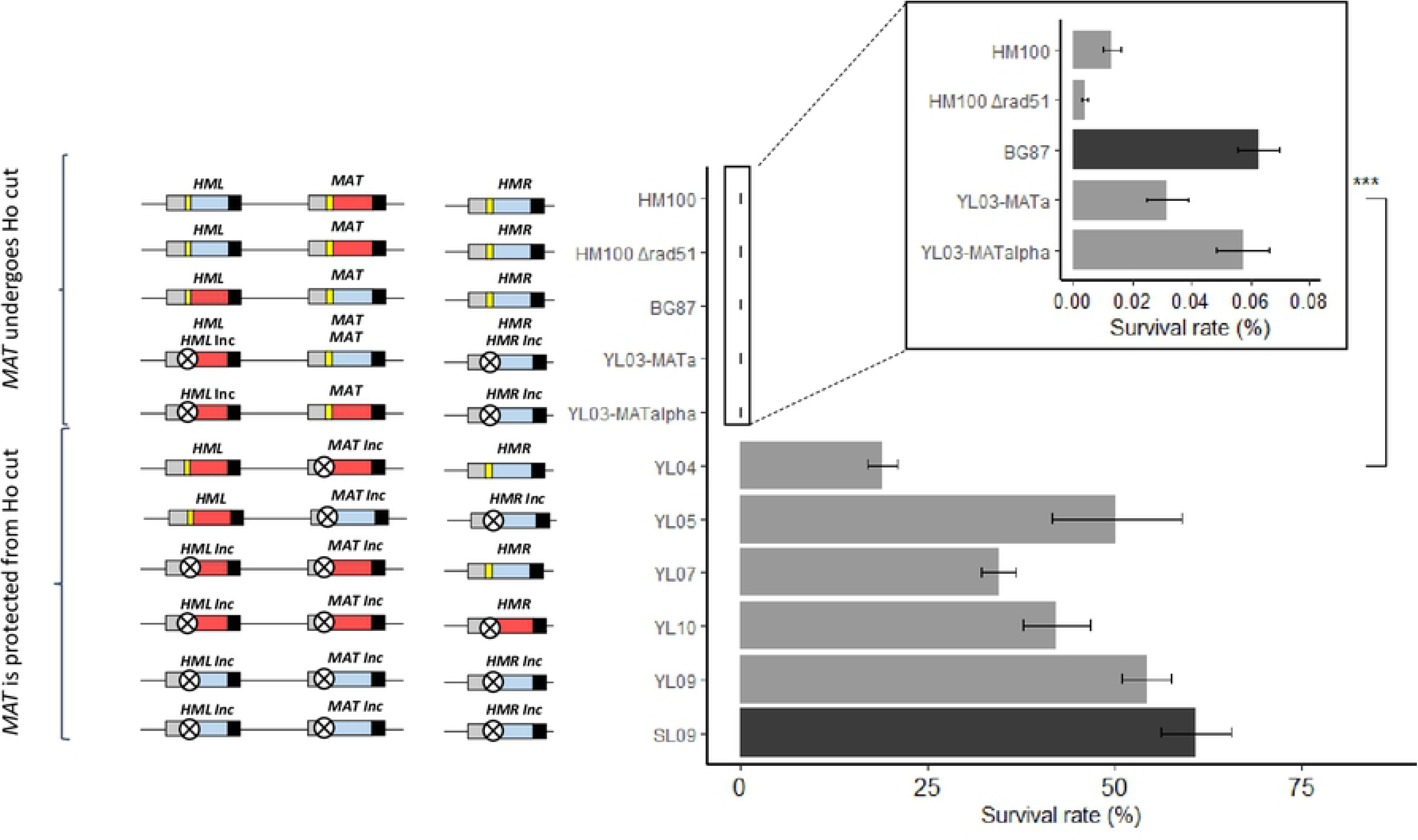
Survival rate of wild-type, *Δrad51* mutant and inconvertible strains upon Ho induction. Blue box represents Ya, red box Yalpha, yellow bar wild-type Ho site and crossed circle mutated Ho site (**inc** loci) (not to scale). Histogram shows survival rate of strains with corresponding *MTL* configuration (black, BG87 background and grey, HM100 background). Results for HM100 and BG87 are from (22). Values from, at least, four experiments were averaged, the SEM used as estimate of the error, and the p-value was calculated using the Wilcoxon test. ***: P-value<0.001.

**Table 2.** Molecular structure of *MTL*s in clones from individual p7.1 transformants

| Strain | Locus screened | Mating-type screened by PCR on tested surviving colonies | Mating-type screened by PCR on tested sub-clones |
| --- | --- | --- | --- |
| HM100 $\Delta rad51$<br>(*HMLalpha MATalpha HMRA*) | *HML*,<br>*MAT* and<br>*HMR* | 39/39 pure *HMLalpha MATalpha HMRA* | ND |
| SL-CG8<br>(*HMLalpha-inc MATa $\Delta hmr$*) | *MAT* | 36/36 pure *MATalpha-inc* | ND |
| SL-CG9<br>($\Delta hml$ *MATa HMRalpha-inc*) | *MAT* | 36/36 pure *MATalpha-inc* | ND |
| YL03-MATa<br>(*HMLalpha-inc MATa HMRA-inc*) | *MAT* | 25/32 pure *MATalpha-inc*<br>6/32 mixed *MATalpha-inc/a-inc*<br>1/32 pure *MATa-inc* | 24/24<br>*MATalpha-inc* |
| YL03-MATalpha<br>(*HMLalpha-inc MATalpha HMRA-inc*) | *MAT* | 42/50 pure *MATalpha-inc*<br>5/50 mixed *MATalpha-inc/a-inc*<br>2/50 pure *MATa-inc*<br>1/50 pure *MATa* | ND |

Surviving colonies obtained on inductive medium are screened by PCR at the locus that can be cut by *Ho* endonuclease. Some mixed colonies are sub-cloned to get the ratio of each mating-type in that kind of colony. ND: Not Done.

### A single DSB at <u>MAT</u> is sufficient to induce cell death in <u>C. glabrata</u>

In order to test whether lethality results from multiple Ho-induced DSBs, we mimicked the situation of *S. cerevisiae* where a single recipient of the Ho-induced DSB, the *MAT* locus can be repaired by the two non-cleavable donors *HML* and *HMR.* We mutated the Ho recognition site of both *HML* and *HMR,* so that only the *MAT* locus can be cut by Ho (strains YL03-MATalpha and YL03-MATa, Table 1). Expression of *HO* in those strains leads to a lethality similar to the one obtained in wild-type strains HM100 and BG87 (Fig 1). Thus, a single Ho-induced DSB at *MAT*, whatever its mating-type, is sufficient to induce massive cell death in *C. glabrata*.

### Reducing the duration of induction of Ho does not save cells from death

We reason that continuous induction of *HO* expression on solid medium for 48 hrs could be lethal due to continuous cutting. To overcome this eventuality, we performed a Ho-induction time course experiment in which Ho is induced in liquid medium and its expression is repressed, at different time points, by plating cells on repressive medium. The survival rate can thus be calculated by the ratio of colonies obtained on repressive medium to the theoretical number of cells plated on this medium. This experiment was done on strain SL-CG9, in which only *MAT* can be cut and repaired by *HMR* (*Δhml MATa HMRalpha-inc*, Table 1), thus allowing us to prevent death issues linked to cutting at *HML* and *HMR*.

As shown on Figure 2A, induction of the Ho-DSB leads to a drastic drop of the survival rate with more than 98 % of cell death at T=2 hrs. The survival rate then keeps slowly decreasing up to T=19 hrs. It then rises, probably because surviving cells invade the liquid culture. Molecular analysis of surviving colonies shows that mating-type switching increases up to T=4 hrs (Fig 2A). These results show that cells cannot be saved by stopping Ho induction, even at early stages of the experiment.

**Fig 2.**
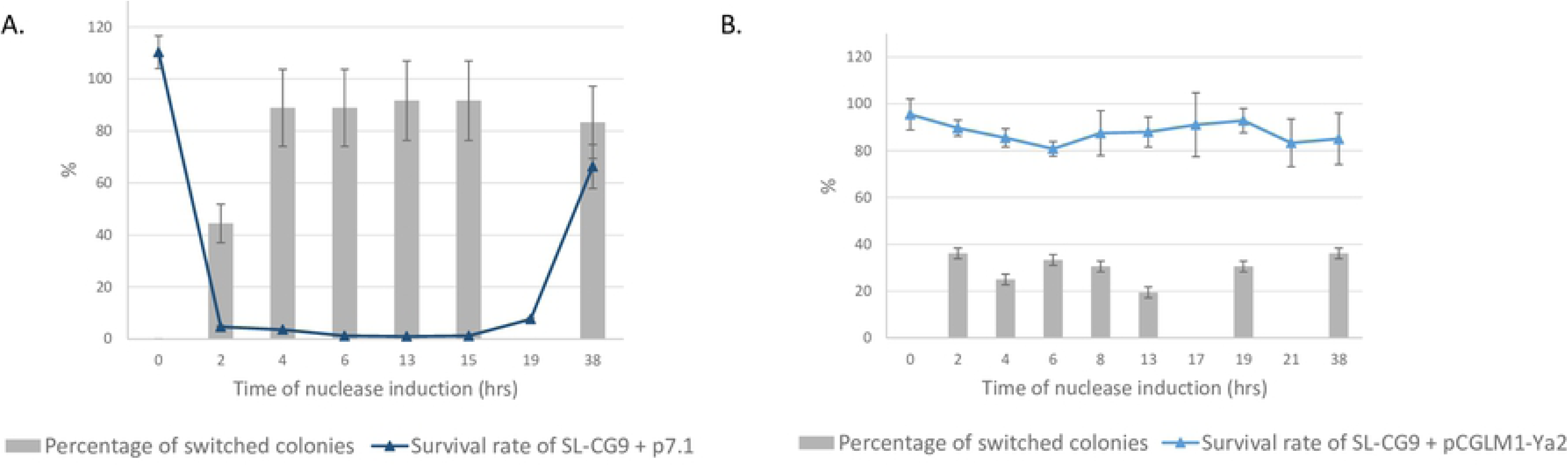
Survival rate during Ho (A) or Cas9 (B) induction associated to the percentage of switched colonies. Induction was performed in liquid during a time-course experiment for strain HM100 expressing Ho (harboring p7.1) (A) or expressing Cas9 targeting *MATa* (harboring pCGLM1-Ya2) (B). The Y-axis represents both the survival rate (curve) expressed as a percentage, and the percentage of switched colonies (histogram). Survival rate is calculated by comparing the number of colony-forming units on SC-Rep with the number of cells plated, as estimated by counting; and is normalized by dividing it by the survival rate of the control strain, i.e. the strain transformed by pCGLM1 for Cas9 induction and the strain transformed by pYR32 for Ho induction, grown in the same conditions. For survival rate, values from four experiments were averaged and the SEM is used as estimate of the error. For the percentage of switched colonies, the square root of the number of surviving colonies screened is used, i.e. sqrt of 36. For time-course experiments, at points T=19 hrs in (A) and T=17 and T=21 hrs in (B), no surviving colonies screen was performed.

### Lethality is not due to toxic recombinational repair intermediates

We wondered whether the fact that *HMR* is not on the same chromosome as *HML* and *MAT*, contrary to *S. cerevisiae*, could be a cause for lethal rearrangements during DSB repair at *MAT*. Alternatively, death could be a result of the degeneration of the mating-type switching mechanism, for example by invasion of both *HML* and *HMR* by the two ends of the broken *MAT* locus, leading to non-resolvable structures.

In order to test this, we constructed two strains in which *MAT* can be cut by Ho and can only be repaired either by *HML* or by *HMR* (SL-CG8, *HMLalpha-inc MATa Δhmr*, and SL-CG9, *Δhml MATa HMRalpha-inc,* respectively, Table 1). Expression of the *HO* gene in both strains leads a high lethality (Fig 3A), similar to the ones of the wild-type or YL03 strains. We analyzed the molecular structure of the *MAT* locus in colonies from induction plates, by PCR using primers specific of the mating-type carried by the *MTL* (“alpha” or “a”, wt or inc, S2 Table and S1 Appendix). This allows the distinction of the original *MAT* locus from the repaired locus that has become resistant to cutting. All surviving colonies tested exhibited mating-type switching, whatever the location of the repair template (*HML* in strain SL-CG8 and *HMR* in strain SL-CG9, Table 2). Thereby, the genomic localization and consequently the configuration of the repair templates in *C. glabrata* is not the cause of lethality.

**Figure 3.**
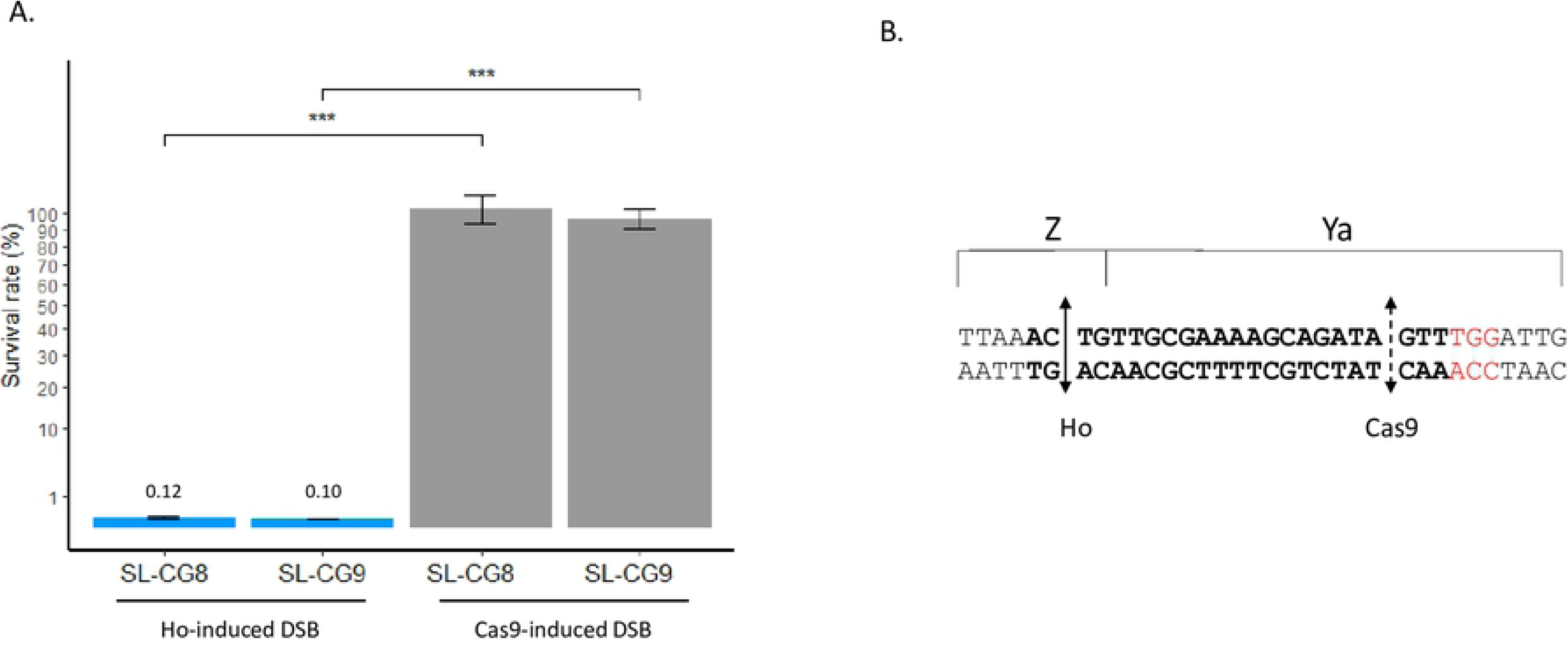
Survival rate upon Ho and Cas9 induction and gRNA used for Cas9. (A) Survival rate of strains SL-CG8 and SL-CG9 upon Ho (in blue) and Cas9 (in grey)-induced DSB at *MAT*. The Y-axis follows a square root scale. Induction is performed on solid medium. Values from four experiments were averaged, the SEM used as estimate of the error, and the P-value was calculated using the Wilcoxon test. ***: P-value<0.001. (B) The gRNA targeting the *MATa* locus of *C. glabrata*. Sequence shown is a segment of the *MATa* locus of HM100, including the gRNA in bold and the PAM sequence in red. Plain double arrow indicates the Ho cleavage site and dashed double arrow the Cas9 cleavage site.

### Donor preference in <u>C. glabrata</u> is biased towards <u>HML</u>

Donor preference in *S. cerevisiae* is a highly regulated mechanism which allows a productive mating-type switching by promoting the use of the donor locus of opposite mating-type to repair the DSB at *MAT* (15). In order to know whether this preference was conserved in *C. glabrata*, we performed a molecular analysis of the *MAT* locus after induction of Ho to reveal which template was used for repair in strains that carry different and inconvertible mating-types at *HML* and *HMR*, i.e.strains YL03-MATalpha and YL03-MATa (Table 1).

Analysis of surviving colonies from strain YL03-MATa shows that 78 % display only the alpha-inc information at *MAT*, 3 % display only the a-inc information at *MAT*, and 19 % show both alpha-inc and a-inc information in the same colony (mixed colonies) (Table 2). The latter can arise if the DSB at *MAT* happens after the first cell division so that cells can repair the DSB, independently, using either *HMLalpha-inc* or *HMRa-inc*. Such mixed colonies were sub-cloned in order to get the ratio of cells that have used *HML* or *HMR* as template but we failed to isolate *MATa-inc* sub-clones, indicating that the use of *HMR* is a very rare event (∼3 %) (Table 2).

Similar results were obtained for strain YL03*-*MATalpha: 84 % of surviving colonies tested display only the *MATalpha-inc* genotype, 4 % display a pure *MATa-inc* genotype, 10 % show both alpha-inc and a-inc information at *MAT* in the same colony and 2 % remain *MATalpha* (Table 2). Thus, in contrast to *S. cerevisiae, HML* is preferentially used as template for repair in *C. glabrata*, whatever the mating-type at *MAT*.

### Protecting the <u>MAT</u> locus from DSB is sufficient to restore viability

We then expressed *HO* in a strain in which only *MAT* is protected from the cut, while both *HML* and *HMR* can be cleaved by Ho (Strain YL04, Table 1). In this strain, cell viability drastically increases to ∼20 %. Survival does not reach 100 % but is 2 000 times higher than in the wild-type isogenic strain, HM100 (P-value<0.001, Wilcoxon test) (Fig 1). This result has been confirmed with five other strains in which *MAT* and either *HML* or *HMR*, or both, are protected from the Ho-induced DSB (Strains YL05 and YL07 respectively, and strains YL10, YL09 and SL09, Table 1 and Fig 1). Thus, the high lethality induced by expression of Ho, disappears when *MAT* is protected from the cut. However, when the three *MTL* loci are protected from the Ho DSB, the survival rate never exceed 61 %. This result underlies a toxic role of *S. cerevisiae*’s Ho outside its role in lethal DSBs at *MAT*.

### The <u>MAT</u>-DSB induced lethality is specific to Ho

We checked whether the *MAT* DSB-induced lethality was caused by the DSB *per se* at *MAT.* In order to test this, we induced a DSB at *MAT* by Cas9 using the CRISPR-Cas9 system from (26). This system relies on a unique *URA3* plasmid, pCGLM1, in which *CAS9* gene is inducible. This allows us to induce a DSB at *MAT* with Cas9, in the same conditions as with Ho in strains SL-CG8 and SL-CG9 (*HMLalpha-inc MATa Δhmr*, and *Δhml MATa HMRalpha-inc*, respectively). We used as gRNA, a sequence that targets the Ho site of the locus containing Ya, so that it can only target the *MAT* locus in strains SL-CG8 and SL-CG9. Due to constraints on the design of the gRNA, the Cas9-induced DSB is shifted by 18 bp compared to the Ho-induced DSB (Fig 3B).

Induction on solid medium shows that 95 % of the cells are able to give rise to a colony, as shown on Figure 3A. Furthermore, the very low lethality observed corresponds to the one observed when *CAS9* is expressed alone (without any gRNA) (26). We made sure that Cas9 had indeed cut the *MAT* locus by screening mating-type switching of surviving colonies by PCR. Depending of the strain, between 87 % and 100 % of the colonies tested presented “alpha-inc” information at *MAT*, even though most are mixed colonies, confirming the cut of this locus by Cas9 and induction of mating-type switching (Table 3).

**Table 3.** Molecular structure of *MAT* in clones from individual pCGLM1-Ya2 transformants

| Strain | Locus screened | Mating-type screened by PCR on tested surviving colonies | Mating-type screened by PCR on tested sub-clones |
| --- | --- | --- | --- |
| SL-CG8<br>(*HMLalpha-incMATa Δhmr*) | *MAT* | 36/40 *MATalpha-inc/a*<br>4/40 *MATalpha-inc* | 40/48 *MATalpha-inc*<br>8/48 *MATa* |
| SL-CG9<br>(*Δhml MATa HMRalpha-inc*) | *MAT* | 32/45 *MATalpha-inc/a*<br>7/45 *MATalpha-inc*<br>6/45 *MATa* | 38/48 *MATalpha-inc*<br>10/48 *MATa* |

Surviving colonies obtained on inductive medium are screened by PCR at the locus that can be cut by the Cas9 endonuclease. Some mixed colonies are sub-cloned to get the ratio of each mating-type in that kind of colonies.

We also performed a time-course experiment with Cas9 in strain SL-CG9 (Fig 2B). This shows that Cas9 induction in liquid medium only leads to a decrease of 20 % of the survival rate at 6 hrs. The survival rate then rises rapidly to become stable after 8 hrs of induction. Surprisingly, contrary to what we observe in induction on plates, screening of mating-type switching at *MAT* revealed that only ∼20 to 36 % of surviving colonies have switched mating-type (Fig 3B).

These results show that the *MAT* locus can be cut and repaired by HR without any accompanying high lethality.

## Discussion

Mating-type switching is a highly regulated mechanism that relies on a chromosomal DSB. DSBs are a major threat for genome integrity (27). Repair of such damage is essential and can be achieved through Rad51-dependent HR which involves many steps in order to succeed: search for homology involving Rad51 and Rad52 in *S. cerevisiae*, copy on the donor locus and displacement and resolution of the double Holliday junction (28). In *S. cerevisiae*, the DSB at the *MAT* locus is repaired by HR using *HMR* or *HML* as template, depending on the original mating-type of the cell. *C. glabrata* does not switch mating types spontaneously at high frequency (20). We have previously shown that mating-type switching can be efficiently induced in this yeast by expressing the *HO* gene from *S. cerevisiae*, but that it is lethal to most cells (22). Our previous work also showed that the *HML* locus is cut in *C. glabrata*; something that never happens in wild-type strains of *S. cerevisiae* (22). In this work, we aimed at understanding the link between mating-type switching and cell death in *C. glabrata*. To this end, we constructed strains with inconvertible Ho sites (**inc**) in which mutations have been introduced precisely on the Ho site in such way that the Ho cut is prevented. We thus could examine survival to individual DSB at the different *MTL* loci as well as knowing which *MTL* has been used as template for repair.

We now show that *HMR* is also cut by the Ho endonuclease in *C. glabrata*, suggesting a deficiency of silencing mechanisms at this locus. This assumption is supported by previous studies that have shown that, in *C. glabrata, HMR* is not silenced at the transcriptional level, and that subtelomeric silencing is less robust than *S. cerevisiae’s* silencing mechanisms (29, 30). In *S. cerevisiae*, the donor preference mechanism ensures an efficient mating-type switching at *MAT* by promoting the use of the template from the opposite mating-type, in repair (15). We found, in *C. glabrata*, that whatever the mating-type at *MAT*, *HML* is preferentially used as template for repair. This indicates that the donor preference from *S. cerevisiae* seems not to be conserved in *C. glabrata* and that the length of the sequence homology shared between the loci, *HML*, *MAT* and *HMR* does not influence the use of the donor for repair of the *MAT* DSB. Along with the fact that the *C. glabrata* endogenous Ho protein fails to induce efficient mating-type switching (22), these results could indicate a degeneration of the mating-type switching system in *C. glabrata.* This cannot be related to the content of *C. glabrata*’s genome as it has retained all the genes known to be involved in DSB repair in *S. cerevisiae* (31). However, it is understandable that such a dangerous mechanism would be lost if it is not essential; as seems to be the case in *C. glabrata* since no sexual cycle has been described in this species.

In our previous work, we hypothesized that multiple DSBs at the *MTL* loci would be unrepairable and that this was the cause for lethality when mating-type switching is induced. To prevent additional DSBs at *HML* and *HMR* and mimic *S. cerevisiae*’s situation, in which *MAT* is the only recipient of the Ho cut, we mutated the Ho site at both *HML* and *HMR*. We are now able to demonstrate that one Ho-DSB at the *MAT* locus is sufficient to induce cell death at a similar level to wild-type cells, thus invalidating our previous hypothesis. This means that, even in the presence of two intact homologous sequences, the *MAT* locus is not able to repair the break. More surprisingly, the DSB at *MAT* is only lethal when it is performed by the Ho protein. We show that mating-type switching can be triggered by CRISPR-Cas9, thus independently of the Ho protein, in *C. glabrata*. This has been shown only recently in the model species *S. cerevisiae* (32). No lethality is observed after a Cas9-DSB at *MAT* on plates and a lethality of ∼20 % is observed in liquid induction. This lethality probably corresponds to the fact that HR is less efficient in *C. glabrata* than in *S. cerevisiae* (33, 34). The discrepancy in survival between solid and liquid induction experiments can be explained by the fact that surviving cells appear early enough on solid induction to give rise to a colony, leading to survival rate of 95 %. The substantial difference between liquid and solid induction resides in the percentage of switched colonies obtained after Cas9 induction. On solid medium, nearly all surviving colonies tested on induction plates showed mating-type switching, suggesting that the Cas9 cut at the *MAT* locus is highly efficient. In addition, this is true whatever the template available for repair, *HML* or *HMR*. It thus demonstrates that both *HML* and *HMR* are accessible repair templates for *MAT,* and that location of *HMR* on another chromosome than *MAT* does not prevent its use as template, nor does it cause lethality. It also shows that the HR system in *C. glabrata* is efficient. On the contrary, only ∼30 % of surviving colonies from Cas9 induction time-course experiment showed a switch at *MAT*, even at 38 hrs of induction. This can be explained by a growth competition in liquid medium between Cas9- and Cas9+ cells. It is probable that cells that have switched mating-types keep a functional plasmid that expresses Cas9 continuously since once they have switched, they become resistant to further cutting. The continuous expression of Cas9 could slow down growth of such cells whereas cells that have mutated the plasmid before switching their mating-type at *MAT* grow faster. Thus, cells that have mutated the *CAS9* gene (or its promoter in such way that *CAS9* is not expressed anymore) invade the liquid culture so that *MAT*-switched cells will be diluted and less represented on repressive plates.

Unless the difference in the lethality between the expression of Cas9 and of Ho is due to the 18 bp shift in cutting, which seems highly unlikely, these results suggest that the Ho protein prevents DSB repair specifically at the *MAT* locus, in such a way that 99.9 % of cells die. This high lethality is specific to *C. glabrata* as expression of *S. cerevisiae*’s *HO* gene, exactly in the same conditions as in *C. glabrata*, is not lethal in the close species *Nakaseomyces delphensis* (unpublished data). It is surprising that *S. cerevisiae*’s Ho could have a deleterious effect in a locus-specific manner. As in all three-loci based mating-type switching systems, the three *MTL* loci of *C. glabrata* share identical sequences and only differ by the mating-type carrying and/or their location in the genome (35). We know that mating-type borne by any of the *MAT* does not influence lethality as both HM100 and BG87 die at 99.99 % (Fig 1). Thus, only the location of the *MAT* locus could explain the specificity of lethality induced by Ho. The *MAT* locus is located in a central region on chromosome B whereas *HML* and *HMR* are positioned in subtelomeric regions on chromosome B and E, respectively (36). Thus, the Ho specificity for *MAT* could only be achieved either through the structure of the chromatin or through the flanking sequences of the *MAT* locus. How exactly does the *S. cerevisiae*’s Ho protein act to induce a high lethality remains unknown but one hypothesis is that Ho prevents DSB repair by getting stuck at *MAT,* after performing the DSB, preventing recruitment of recombination proteins and thus repair of the locus.

Finally, we would like to discuss the toxic role of *S. cerevisiae*’s Ho, in *C. glabrata*, outside its role in lethal DSB at *MAT*. Several hypotheses may be envisaged. In *C. glabrata*, *S. cerevisiae*’s Ho endonuclease could cut another site in the genome, outside the three *MTL* loci that would be lethal in a haploid genome. Even if, by a bioinformatics analysis, we could not find any additional Ho sites outside the *MTL* loci, we cannot exclude the existence of a more degenerate site. An alternative hypothesis is that the Ho protein binds the mutated Ho sites (**inc**) and gets stuck there. In that way, it could, for example, physically block replication forks and thus prevent DNA replication and cell division. We favor the second hypothesis as in a strain in which *MAT* is deleted and both *HML* and *HMR* are inconvertible, survival rate reaches ∼83 % (unpublished data). Performing a ChIP-PCR on the Ho protein to examine its binding on the three *MTL* loci would allow us to better explore this aspect.

## Materials and methods

### Strain, cultures and transformation

*C. glabrata* strains used in this study are listed in Table 1. Strains are grown in broth or on plates at 28°C in YDP (non-selective, 1% Yeast Extract, 1% Peptone, 2% glucose), in Synthetic Complete medium lacking uracil (SC-Ura, 0.34 % Yeast Nitrogen Base without amino acids, 0.7 % ammonium sulfate, 2 % glucose, supplemented with adenine and all amino acids except uracil) or in Synthetic Complete medium lacking uracil, methionine, and cysteine (induction conditions for the *MET3* promoter, SC-Ind, 0.34 % Yeast Nitrogen Base without amino acids, 0.7 % ammonium sulfate, 2 % glucose, supplemented with adenine and all amino acids except methionine and cysteine). For selection of transformants of the Ho plasmid or Cas9 plasmid and maintenance in repressive conditions for the *MET3* promoter, strains are grown in SC-Ind supplemented with 2 mM each of methionine and cysteine (SC-rep) and in YPD supplemented with 2 mM each of methionine and cysteine (YDP-Rep) when repression but no selection is needed. For SC-Rep, medium is buffered by 10 mL of Na_2_HPO_4_ 0.05 M and NaH_2_PO_4_ 0.95 M per liter. For *URA3* counter-selection marker, yeast strains are grown on 5-FOA medium (SC-Ura supplemented with 1 g/L of 5-fluoroorotic acid (5-FOA) and 50 mg/L of uracil).

Transformation is done according to the “one-step” lithium acetate transformation protocol from (36).

### Induction of mating-type switching by Ho

The *HO* gene from *S. cerevisiae* is cloned into the pCU-MET3 plasmid under the *MET3* promoter (p7.1, S1 Table) (37) and protocol for solid induction is detailed in (22). For time-course of induction in liquid medium, transformants are grown overnight in liquid SC-Rep medium, counted, washed and resuspended in sterile water at 4.10^7^ cells/mL. 100 µL is used to inoculate 40 mL of liquid SC-Ind medium and the culture is placed at 28°C with agitation. For each time point, a sample of the culture is counted under the microscope, diluted and plated on SC-Rep plates.

### Induction of mating-type switching by CRISPR-Cas9

We used the inducible CRISPR-Cas9 system for *C. glabrata* from (26) through plasmid pCGLM1. We cloned into pCGLM1 a sequence corresponding to a guide RNA (gRNA) targeting the Ya sequence (S2 Table), giving rise to plasmid pCGLM1-Ya2. Induction of Cas9 DSB was then performed as for induction of the *S. cerevisiae*’s *HO* gene done with p7.1 (see above).

### Construction of strains

We mutated the Ho sites in the region known to be essential for Ho cutting in *S. cerevisiae* (38), as shown on Appendix 2, yielding loci *HML-inc MAT-inc* and *HMR-inc.* Modification of *HML*, *MAT*, or *HMR* loci was realized either by marker selection (pop-in/pop-out) (39) or by mating-type switching upon *HO* gene expression or by use of CRISPR/Cas9. The three methods are detailed below. Primers and plasmids are listed in S2 and S1 Tables, respectively. Method used to construct each strain is listed in S3 Table.

#### Construction of PCR fragments and plasmids for pop-in

In order to integrate the *URA3* marker at the targeted locus (pop-in), we amplify the *URA3* gene from *S. cerevisiae* under its own promoter by PCR using primers Sc-URA3-F and Sc-URA3-R and, YEp352 as template. The PCR fragment is cloned into the *EcoR*V-digested pBlueScript. Such cloning gives rise to pURA (S1 Table).

To direct integration of the *URA3* marker at the targeted locus, here the *MTL* loci *HML*, *MAT* or *HMR,* the 5’ and 3’ flanking regions are added to the *URA3* marker in multiple steps. First, the Z sequence, shared by the three *MTL* loci, was amplified by PCR using primers 68/70 and HM100 strain DNA as template (S2 Table). Primers 68 and 70 contain *BamH*I and *EcoR*I restriction sites, respectively, to allow cloning of the Z PCR fragment upstream of the *URA3* marker into pURA, giving rise to pZUA (S1 Table).

Second, Ya and Yalpha sequences were amplified on strain HM100 by PCR, using primers 73/72 and 74/69 respectively (S2 Table). Primers 73 and 72 contain *Hind*III and *Sal*I restriction sites, respectively, in order to clone the Ya PCR fragment downstream of the *URA3* marker into pZU, giving rise to pZUA (S1 Table). The *Sal*I restriction site was added to primer 69 and no restriction site was added to primer 74 as the Yalpha PCR fragment already contains the *Hind*III restriction site 38 bp from the 5’ of the fragment. Thus, the Yalpha PCR fragment, digested by both *Sal*I and *Hind*III, was cloned downstream of the *URA3* marker into pZU to give rise to pZUAlpha (S1 Table).

Amplification by PCR, using universal primers M13F/M13R, on the both pZUA and pZUAlpha plasmids, leads to ZUA and ZUAlpha fragments, respectively. These fragments have been used for targeting *HML*, MAT or *HMR* loci (S1 Table) and Ura+ transformants were selected on SC-Ura. Correct integration of the fragment was checked by PCR.

#### Construction of plasmids and PCR fragments for pop-out

The *URA3* marker is removed (pop-out) from the target locus by homologous recombination with a DNA fragment derived from the upstream and downstream sequences of that locus (S3 Table).

In order to replace the wild-type Ho site in the different *MTL* loci, by the inconvertible-mutated Ho site, we constructed two plasmids; pZA-inc and pZalpha-inc (S1 Table). The pZA-inc plasmid (without *URA3* gene) results from double digestion of pZUA by *EcoRI* and *HindIII* and ligation after Klenow fill-in. The pZAlpha-inc (without the *URA3* gene) plasmid was constructed by cloning the *BamH*I/*EcoR*I-digested Z fragment and the *EcoR*I/*Sal*I-digested Yalpha fragment into the pBlueScript double digested by *BamH*I and *Sal*I. Amplification by PCR using primers M13F/M13R, from both pZA-inc and pZAlpha-inc plasmids, lead to the ZA-inc and ZAlpha-inc fragments that have been used for pop-out. The wild-type and inconvertible Ho site sequences comparison is presented on Fig. S1. In the case of the fragment used for pop-out of *URA3* for the deletion of *HMR* in strain BG87, construction was done by amplification of upstream and downstream sequences (500 bp each) of *HMR* on strain BG87, using primers Up-HMR-F/Up-HMR-R and Down-HMR-F/ Down-HMR-R, respectively (S2 Table). Primer Up HMR-R contains 40 bp of homology to the 5’ end of the downstream PCR fragment. These two fragments were then combined by fusion PCR using primers Up-HMR-F and Down-HMR-R, giving rise to the ΔHMR fragment (strain SL01, S3 Table). As shown in S3 Table, other fragments for pop-out experiments were obtained by direct PCR on genomic DNA.

About 1 µg of each pop-out fragment was used to transform Ura+ strains, which were then plated onto YPD, grown for 24 hrs and replica-plated onto 5-FOA plates. Resulting 5-FOA^R^ colonies were checked by PCR for correct removal of the *URA3* marker, and the locus has been sequenced in the final strains.

#### Strains obtained by mating-type switching

When possible, we took advantage of the efficient mating-type switching induced by expression of *HO* to transpose the **inc**-Ho site mutation from a sexual locus to another, instead of doing pop-in/pop-out transformations as above. For example, an *HMLalpha-inc* locus can easily be used as template, during gene conversion, to repair either *MAT* wt or *HMR* wt. In addition, extra-chromosomal copies of either *MATa-inc* or *MATalpha-inc* were also used as templates for mating-type switching of *MTL* loci, in order to insert **inc**-*Ho* sites. These copies were introduced in the p7.1 plasmid, as follows. Plasmid p7.1 (22) was digested by K*pn*I, and *MATa-inc* and *MATalpha-inc* sequences were amplified by PCR using primers Up-Rec-MAT-F/ Down-Rec-MAT-R on strains YL07 and YL09, respectively (Table 1 and S2). Both primers share, respectively, 40 bp of homology to the ends of the *Kpn*I-digested plasmid. This allows PCR fragment cloning in p7.1, at the *Kpn*I restriction site, by homologous recombination in *E. coli* (40). Correct assembly was confirmed by both analytic colony PCR and restriction digests.

Expression of Ho is induced in strains that are targeted for modification, either from the p7.1 plasmid, when a genomic *MTL* locus is used as template, or from p7.1-derived plasmids that contain a copy of *MATa-inc* or *MATalpha-inc*. Final loci are checked by PCR and sequencing.

#### Construction of the <u>Δrad51</u> mutant using CRISPR-Cas9

The *Δrad51* mutant of strain HM100 was constructed with the CRISPR-Cas9 system on plasmid pJH-2972 (kind donation from J. Haber, https://protocolexchange.researchsquare.com/article/nprot-5791/v1). We cloned a sequence corresponding to a guide RNA (gRNA) targeting the *RAD51* gene into plasmid pJH-2972 (S2 Table), giving rise to plasmid pJH-RAD51.

We amplified upstream and downstream sequences (500 bp each) of *RAD51* CDS on strain HM100 by PCR using primers Up-Rad51-F/Up-Rad51-R and Down-Rad51-F/Down-Rad51-R, respectively (S2 Table). Primer Up-Rad51-R contains 40 bp of homology to the 5’ end of the downstream PCR fragment. These two fragments are then combined by fusion PCR using primers Up-Rad51-F and Down-Rad51-R, giving rise to the *Δrad51* fragment.

The strain was then co-transformed with both 1 µg of pJH-RAD51 and 1 µg of *Δrad51* fragment. Ura+ transformants were then selected on SC-Ura and checked for deletion at the *RAD51* locus by PCR. Deletion was confirmed by sequencing the *RAD51* locus and by Southern blot analysis (S3 Appendix).

### Cell viability estimation

Different dilutions of cultures, containing between 200 to 10^6^ cells, are spread on both inductive and repressive media. When the survival rate is over 20 %, cell viability is determined directly as the ratio of the number of colonies counted on inductive medium to the number of colonies counted on repressive medium, for the same dilution. When the survival rate is under 1 %, colonies are confluent on repressive medium at the same dilution where several colonies can be observed on induction medium. Thus, survival rate is measured by first comparing the number of colony-forming units (CFU) on inductive medium with the theoretical number of cells plated, as estimated by counting on a Thoma counting chamber. This is then corrected by the ratio of CFU to the number of cells counted, estimated by plating 200 cells on repressive medium. All the values were obtained from at least four independent transformants. Colonies number from a minimum of 18 to a maximum 494 was counted on plates.

### Determining the genotype at *MTL* loci

The genotype of surviving colonies at each *MTL* locus is determined by PCR using specific primers: the forward primer is located upstream of the locus (ensuring specificity of the locus screened; *HML*, *MAT* or *HMR*) and a reverse primer located precisely on the Ho site (ensuring specificity of the information carried by the locus; alpha or a and wt or **inc**) (S2 Table, S1 Appendix). We checked that the mutated Ho-sites, “alpha-**inc**” and “a-**inc**” are not cut since we never observe switching at those loci (not shown).

## Acknowledgements

We thank members of our lab and the iGénolevures network ((IRN from the CNRS N°0814) for stimulating discussions, Gilles Fischer, Fabienne Malagnac and Pierre Grognet for critical reading.

## Supporting information captions

**S1 Appendix. Mating-type screened at *MAT* in different strains**

All strains are analyzed with primer pairs that are specific to *MATa*, *MATalpha*, *MATa*-inc and *MATalpha*-inc, respectively GS01/123, GS01/121, GS01/122 and GS01/120. Top left panel: amplification obtained on BG87 (*MATa*); bottom left panel: amplification obtained on YL05 (*MATa*-inc); top right panel: amplification obtained on HM100 (*MATalpha*); bottom right panel: amplification obtained on YL07 (*MATalpha*-inc). MM: Molecular Marker, GeneRuler 1 kb (Thermo Fisher Scientific Inc).

**S2 Appendix. Comparison of wild-type and mutated Ho site of locus carrying Yalpha (A) or Ya information (B)**.

The wild-type Ho site is shown on top in blue letters, the mutated Ho site is shown below with mutated bp in red and deleted bp as dashes. Arrows indicate the Ho cleavage site.

**S3 Appendix. Molecular characterization of knock-out HM100 *Δrad51* mutants by Southern blot hybridization**.

Two restriction enzyme digestions have been performed to confirm the correct deletion of *RAD51* in strain HM100: one with *Hind*III (left) and one with *Nde*I/*EcoR*I (right). In both cases, the same probe has been used, corresponding to 500 bp homologous to the 5’ UTR fused to 500 bp homologous to the 3’ UTR.

**S1 Table.** Plasmids used in this work.

**S2 Table.** Primers used in this work. Fw: Forward; Rv: Reverse. The lowercase letters represent sequence with no homology to template DNA, whereas homologous regions are indicated in uppercase.

**S3 Table.** Methods used for strains construction.

## Bibliography

1. Ni M, Feretzaki M, Sun S, Wang X, Heitman J. Sex in fungi. Annu Rev Genet. 2011;45:405–30.

2. Butler G. Fungal sex and pathogenesis. Clin Microbiol Rev. janv 2010;23(1):140–59.

3. Heitman J, Carter DA, Dyer PS, Soll DR. Sexual reproduction of human fungal pathogens. Cold Spring Harb Perspect Med. 1 août 2014;4(8).

4. Bennett RJ, Johnson AD. Completion of a parasexual cycle in Candida albicans by induced chromosome loss in tetraploid strains. EMBO J. 15 mai 2003;22(10):2505–15.

5. O’Gorman CM, Fuller HT, Dyer PS. Discovery of a sexual cycle in the opportunistic fungal pathogen Aspergillus fumigatus. Nature. 22 janv 2009;457(7228):471–4.

6. Hanson SJ, Wolfe KH. An Evolutionary Perspective on Yeast Mating-Type Switching. Genetics. 2017;206(1):9–32.

7. Egel R. Fission yeast mating-type switching: programmed damage and repair. DNA Repair (Amst). 2 mai 2005;4(5):525–36.

8. Maki T, Ogura N, Haber JE, Iwasaki H, Thon G. New insights into donor directionality of mating-type switching in Schizosaccharomyces pombe. PLoS Genet [Internet]. 31 mai 2018 [cité 17 juill 2018];14(5). Disponible sur: https://www.ncbi.nlm.nih.gov/pmc/articles/PMC6007933/

9. Barsoum E, Martinez P, Aström SU. Alpha3, a transposable element that promotes host sexual reproduction. Genes Dev. 1 janv 2010;24(1):33–44.

10. Lee C-S, Haber JE. Mating-type Gene Switching in Saccharomyces cerevisiae. Microbiol Spectr. avr 2015;3(2):MDNA3-0013–2014.

11. Haber JE. Mating-type genes and MAT switching in Saccharomyces cerevisiae. Genetics. mai 2012;191(1):33–64.

12. Tomimatsu N, Mukherjee B, Harris JL, Boffo FL, Hardebeck MC, Potts PR, et al. DNA-damage-induced degradation of EXO1 exonuclease limits DNA end resection to ensure accurate DNA repair. J Biol Chem. 30 juin 2017;292(26):10779–90.

13. White CI, Haber JE. Intermediates of recombination during mating type switching in Saccharomyces cerevisiae. EMBO J. mars 1990;9(3):663–73.

14. Loo S, Rine J. Silencers and domains of generalized repression. Science. 17 juin 1994;264(5166):1768–71.

15. Herskowitz I. Life cycle of the budding yeast Saccharomyces cerevisiae. Microbiol Rev. déc 1988;52(4):536–53.

16. Dujon B, Sherman D, Fischer G, Durrens P, Casaregola S, Lafontaine I, et al. Genome evolution in yeasts. Nature. 1 juill 2004;430(6995):35–44.

17. Muller H, Hennequin C, Gallaud J, Dujon B, Fairhead C. The asexual yeast Candida glabrata maintains distinct a and alpha haploid mating types. Eukaryotic Cell. mai 2008;7(5):848–58.

18. Ramírez-Zavaleta CY, Salas-Delgado GE, De Las Peñas A, Castaño I. Subtelomeric silencing of the MTL3 locus of Candida glabrata requires yKu70, yKu80, and Rif1 proteins. Eukaryotic Cell. oct 2010;9(10):1602–11.

19. Robledo-Márquez K, Gutiérrez-Escobedo G, Yáñez-Carrillo P, Vidal-Aguiar Y, Briones-Martín-Del-Campo M, Orta-Zavalza E, et al. Candida glabrata encodes a longer variant of the mating type (MAT) alpha2 gene in the mating type-like MTL3 locus, which can form homodimers. FEMS Yeast Res. 2016;16(7).

20. Butler G, Kenny C, Fagan A, Kurischko C, Gaillardin C, Wolfe KH. Evolution of the MAT locus and its Ho endonuclease in yeast species. Proc Natl Acad Sci USA. 10 févr 2004;101(6):1632–7.

21. Carreté L, Ksiezopolska E, Pegueroles C, Gómez-Molero E, Saus E, Iraola-Guzmán S, et al. Patterns of Genomic Variation in the Opportunistic Pathogen Candida glabrata Suggest the Existence of Mating and a Secondary Association with Humans. Curr Biol. 8 janv 2018;28(1):15–27.e7.

22. Boisnard S, Zhou Li Y, Arnaise S, Sequeira G, Raffoux X, Enache-Angoulvant A, et al. Efficient Mating-Type Switching in Candida glabrata Induces Cell Death. PLoS ONE. 2015;10(10):e0140990.

23. Gabaldón T, Martin T, Marcet-Houben M, Durrens P, Bolotin-Fukuhara M, Lespinet O, et al. Comparative genomics of emerging pathogens in the Candida glabrata clade. BMC Genomics. 14 sept 2013;14:623.

24. Fidel PL, Cutright JL, Tait L, Sobel JD. A murine model of Candida glabrata vaginitis. J Infect Dis. févr 1996;173(2):425–31.

25. Cormack BP, Falkow S. Efficient homologous and illegitimate recombination in the opportunistic yeast pathogen Candida glabrata. Genetics. mars 1999;151(3):979–87.

26. Maroc L, Fairhead C. A new inducible CRISPR-Cas9 system useful for genome editing and study of double-strand break repair in Candida glabrata. Yeast. 18 août 2019;

27. Fairhead C, Dujon B. Consequences of unique double-stranded breaks in yeast chromosomes: death or homozygosis. Molec Gen Genet. août 1993;240(2):170–80.

28. Haber JE. DNA repair: the search for homology. Bioessays. mai 2018;40(5):e1700229.

29. Muller H, Hennequin C, Gallaud J, Dujon B, Fairhead C. The Asexual Yeast *Candida glabrata* Maintains Distinct **a** and α Haploid Mating Types. Eukaryotic Cell. mai 2008;7(5):848–58.

30. Ramírez-Zavaleta CY, Salas-Delgado GE, De Las Peñas A, Castaño I. Subtelomeric Silencing of the *MTL3* Locus of Candida glabrata Requires yKu70, yKu80, and Rif1 Proteins. Eukaryotic Cell. oct 2010;9(10):1602–11.

31. Richard G-F, Kerrest A, Lafontaine I, Dujon B. Comparative genomics of hemiascomycete yeasts: genes involved in DNA replication, repair, and recombination. Mol Biol Evol. avr 2005;22(4):1011–23.

32. Xie Z-X, Mitchell LA, Liu H-M, Li B-Z, Liu D, Agmon N, et al. Rapid and Efficient CRISPR/Cas9-Based Mating-Type Switching of Saccharomyces cerevisiae. G3 (Bethesda). 04 2018;8(1):173–83.

33. Cormack BP, Falkow S. Efficient homologous and illegitimate recombination in the opportunistic yeast pathogen Candida glabrata. Genetics. mars 1999;151(3):979–87.

34. Corrigan MW, Kerwin-Iosue CL, Kuczmarski AS, Amin KB, Wykoff DD. The fate of linear DNA in Saccharomyces cerevisiae and Candida glabrata: the role of homologous and non-homologous end joining. PLoS ONE. 2013;8(7):e69628.

35. Muller H, Hennequin C, Dujon B, Fairhead C. Ascomycetes: the Candida MAT Locus: Comparing MAT in the Genomes of Hemiascomycetous Yeasts. In: Taylor JW, Kronstad JW, Heitman J, Casselton LA, editors Sex in Fungi. 1re éd. American Society for Microbiology; 2007. p. 247–263.

36. Gietz RD, Schiestl RH, Willems AR, Woods RA. Studies on the transformation of intact yeast cells by the LiAc/SS-DNA/PEG procedure. Yeast. 15 avr 1995;11(4):355–60.

37. Zordan RE, Ren Y, Pan S-J, Rotondo G, De Las Peñas A, Iluore J, et al. Expression plasmids for use in Candida glabrata. G3 (Bethesda). 3 oct 2013;3(10):1675–86.

38. Nickoloff JA, Chen EY, Heffron F. A 24-base-pair DNA sequence from the MAT locus stimulates intergenic recombination in yeast. Proc Natl Acad Sci USA. oct 1986;83(20):7831–5.

39. Alani E, Cao L, Kleckner N. A method for gene disruption that allows repeated use of URA3 selection in the construction of multiply disrupted yeast strains. Genetics. août 1987;116(4):541–5.

40. Beyer HM, Gonschorek P, Samodelov SL, Meier M, Weber W, Zurbriggen MD. AQUA Cloning: A Versatile and Simple Enzyme-Free Cloning Approach. PLOS ONE. 11 sept 2015;10(9):e0137652.

